# Late stages of the Zika virus life cycle are impaired by a selective TRPML2 agonist

**DOI:** 10.1101/2024.02.01.578205

**Authors:** Kerstin K. Schwickert, Mirco Glitscher, Daniela Bender, Robin Murra, Kevin Schwickert, Steffen Pfalzgraf, Tanja Schirmeister, Ute A. Hellmich, Eberhard Hildt

## Abstract

The flavivirus genus includes human pathogenic viruses such as Dengue (DENV), West Nile (WNV) and Zika virus (ZIKV) posing a global health threat due to limited treatment options. Ion channels are crucial for various viral life cycle stages, but their potential as targets for antivirals is often not fully realized due to the lack of selective modulators. Here, we observe that the human endolysosomal cation channel TRPML2 agonist ML2-SA1 impairs the late life cycle stages of ZIKV, thus underscoring TRPML2 as a promising antiviral target. Upon treatment with ML2-SA1, levels of intracellular genomes and number of released virus particles of two different ZIKV isolates were significantly reduced. ML2-SA1-treated cells displayed enlarged vesicular structures and multivesicular bodies with ZIKV envelope protein accumulation. However, no increased ZIKV degradation in lysosomal compartments was observed. Rather, the antiviral effect of ML2-SA1 seemed to manifest by the compound’s negative impact on genome replication. Moreover, ML2-SA1 treatment also led to intracellular cholesterol accumulation. ZIKV as well as many other viruses including the Orthohepevirus Hepatitis E virus (HEV) rely on the endolysosomal system and are affected by intracellular cholesterol levels to complete their life cycle. Since we observed ML2-SA1 to also negatively impact HEV infections in vitro, this compound may harbor a broader antiviral potential through perturbing the intracellular cholesterol distribution. Besides underscoring the potential of TRPML2 as a promising target for combatting viral infections, we uncover a tentative connection between this protein and cholesterol distribution within the context of infectious diseases.

## 1 Introduction

Zika virus (ZIKV) is a member of the *Flaviviridae* family and gained public attention during a major outbreak in the Americas in 2015/2016 when the infection was first associated with severe neurological complications, such as neonatal microcephaly (Schuler-Faccini et al., 2016). Until now, no specific treatment or prophylactic vaccine is available. ZIKV uses clathrin-mediated endocytosis to enter host cells (Agrelli et al., 2019). It is assumed that subsequent exposure to the acidic environment of the endolysosomal system triggers conformational changes of the ZIKV envelope (E) glycoprotein, which leads to membrane fusion and release of the viral RNA (Dai et al., 2016; Nour et al., 2013). Due to their involvement in viral trafficking and release, endolysosomal ion channels have garnered increasing attention in infectious diseases and as potential antiviral targets (Chao et al., 2023; Grimm et al., 2017; Spix et al., 2020).

The three members of the human transient receptor potential mucolipin (TRPML) subfamily of ion channels mediate the pH-dependent cation release from the endolysosomal system and are involved in membrane trafficking and autophagy (Goretzki et al., 2021; Kim et al., 2009; Li et al., 2017; Sun et al., 2000; Vergarajauregui et al., 2008; Viet et al., 2019; Zeevi et al., 2009). TRPML2 is mainly expressed in lymphocytes and other cells of the immune system (García-Añoveros and Wiwatpanit, 2014). The expression of TRPML2 in mouse macrophages is strongly upregulated after Toll-like receptor activation and TRPML2 plays a role in chemokine and cytokine secretion of macrophages (Plesch et al., 2018; Sun et al., 2015). TRPML2 was reported as a putative interferon-stimulated gene and a potential function of TRPML2 in the innate immune response and during viral infections has been suggested (Lanford et al., 2006; Rinkenberger and Schoggins, 2018; Schoggins, 2019; Spix et al., 2020). Overexpression of TRPML2 in A549 cells was shown to increase infection with endocytosed RNA virus families including *Flaviviridae* (e.g. ZIKV) and *Orthomyxoviridae* (e.g. Influenza A virus (IAV)) by enhancing viral trafficking efficiency through the endolysosomal system (Rinkenberger and Schoggins, 2018). A knockout of TRPML2 decreased infection with IAV (Rinkenberger and Schoggins, 2018). Together, these data indicate that TRPML2 may be a promising antiviral target.

Several synthetic TRPML agonists are available (Chen et al., 2014; Grimm et al., 2012; Plesch et al., 2018; Shen et al., 2012). Their application to cells increases channel-mediated Ca^2+^ flux from endosomes and lysosomes into the cytosol, thereby influencing vesicular fusion and fission, vesicular trafficking, lysosomal exocytosis and autophagy (Medina et al., 2015; Samie et al., 2013). Treatment with the unselective TRPML channel agonist ML-SA1 reduced DENV2 and ZIKV mRNA, protein and the amount of released infectious particles in a dose-dependent manner in A549 and Huh7 cells (Xia et al., 2020). The authors suggested that viral entry, but not the later stages of the DENV2 and ZIKV life cycles were affected by ML-SA1 (Xia et al., 2020). In contrast, a shared TRPML1 and TRPML3 agonist, MK6-83 (Chen et al., 2014), did not display antiviral potential, while the selective TRPML3 agonist SN-2 provoked an antiviral effect (Xia et al., 2020). It was postulated that the antiviral activity of ML-SA1 and SN-2 may be related to effects on TRPML2 and TRPML3 channel activity and expression levels (Xia et al., 2021). However, to date it remains unclear which of the three human TRPML channels are important for different stages during viral infection, and whether all three proteins are equally suited as antiviral targets.

For TRPML2, a potent selective agonist, ML2-SA1 is available (Plesch et al., 2018) that allowed us to investigate the specific role of TRPML2 for flaviviral infections *in vitro*. Two ZIKV isolates, a representative of the Asian lineage (ZIKV French Polynesia) and a representative of the African lineage (ZIKV Uganda), were used. We found that independent of the viral lineage, ML2-SA1 treatment impaired the late stages of the ZIKV life cycle. This suggests a role for TRPML2 in flaviviral replication. We also observed a redistribution of intracellular cholesterol in ML2-SA1-treated cells. Notably, cholesterol-accumulation controls the release pathway of Hepatitis E virus (HEV) negatively, resulting in a lysosomal degradation of capsid structures (Glitscher et al., 2021b) explaining the antiviral effect of ML2-SA1 that we observed against this virus. All in all, this underlines the suitability of TRPML2 as a target for antivirals against ZIKV as well as HEV, and the selective endolysosomal channel agonist ML2-SA1 as a promising lead for novel antiviral compounds.

## 2 Materials & Methods

### 2.1 Cells and viruses

Human epithelial lung carcinoma cells (A549, ATCC CCL-185) and African green monkey kidney (Vero, ATCC CCL-81) cells were maintained in Dulbecco’s modified Eagle medium (DMEM) containing 4.5 g/L D-glucose (Sigma Aldrich, Bio&Sell) supplemented with 10% fetal bovine serum (FBS, Bio&Sell), 100 U/mL penicillin and 100 µg/mL streptomycin (Paul-Ehrlich-Institute central laboratory) in 5% CO_2_ and 95% relative humidity at 37 °C. For ZIKV infection experiments, cells were infected with either the ZIKV French Polynesia (H/PF/2013) or the Uganda (976 Uganda) isolate (both purchased from the European Virus Archive) with a multiplicity of infection (MOI) of 1. Unless otherwise stated, the inoculum was removed 16 h post infection and cells were washed once with PBS (phosphate buffered saline). At the desired time points, supernatants were collected and stored at −80 °C (where applicable) and cells were washed with PBS and lysed for subsequent experiments. Treatment with compounds was started two hours before infection and substances were refreshed during infection and 16 h and 24 h post infection to guarantee constant levels of compounds in cells. For HEV infection experiments, persistently HEV-infected A549 cells (parental A549 cells initially infected with HEV genotype 3c isolate 47832c (Johne et al., 2014)) were used and cells were treated with ML2-SA1 for 24 h.

### 2.2 TRPML agonists and other compounds

The non-specific TRPML agonist ML-SA1 was purchased from Merck KGaA. The selective TRPML2 agonist ML2-SA1 was synthesized according to a two-step procedure from McIntosh *et al*. (McIntosh et al., 2012) and Plesch *et al*. (Plesch et al., 2018). Starting from 2,6-dichlorobenzaldehyde, the benzaldoxime ***(2)*** was prepared by treatment with hydroxylamine hydrochloride. In the second step, the benzaldoxime ***(2)*** was brought to reaction with norbornene in the presence of (bis(trifluoroacetoxy)iodo)benzene (PIFA) to yield ML2-SA1 as a mixture of two enantiomers (for details see SI and Fig. S1–S4). Bafilomycin A1 was purchased from Sigma-Aldrich, Ribavirin from Selleckchem and Leupeptin from AppliChem GmbH. All compound stocks for biological assays were prepared in dimethyl sulfoxide (DMSO) except for Leupeptin, which was dissolved in H_2_O.

**Scheme 1:**
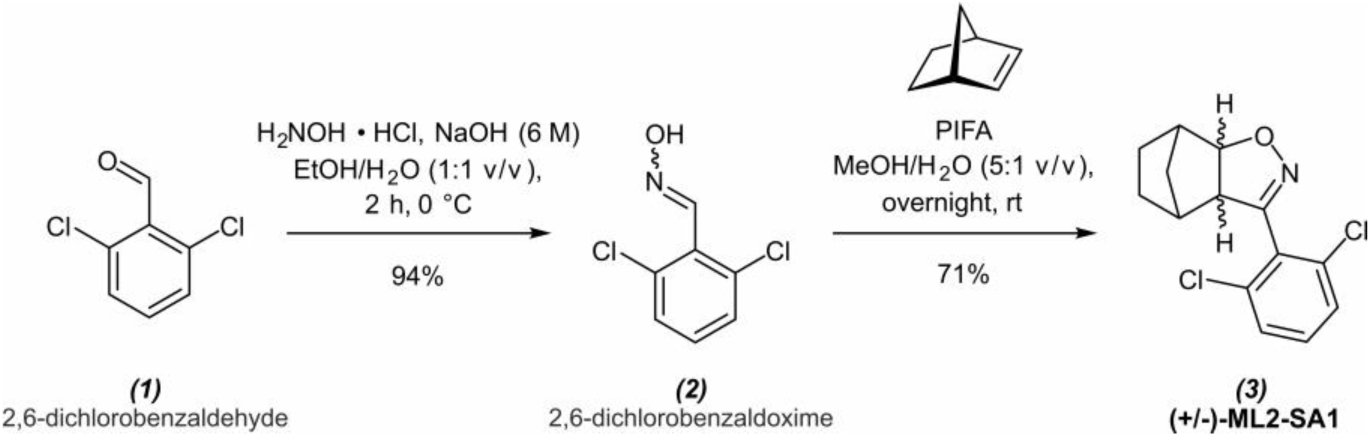
Synthesis of the specific TRPML2 agonist ML2-SA1 following the protocols by McIntosh *et al*. (McIntosh et al., 2012) and Plesch *et al*. (Plesch et al., 2018). For analysis, see Fig. S1–S4.

### 2.3 Cell viability assay

Cell viability assays were performed using the PrestoBlue® Cell viability reagent (Thermo Fisher Scientific) according to the manufacturer’s protocol. A549 cells were seeded at a density of 1×10^4^ cells/well in a flat-bottom 96-well plate and treated with ML2-SA1 and ML-SA1 dissolved in DMSO, final concentrations ranging from 12.5 µM to 200 µM for 24 h and 48 h, respectively. DMSO concentration in all wells was kept constant at 1% v/v. After an incubation period of 1 h at 37 °C with PrestoBlue® Cell viability reagent, the fluorescence of the reagent was measured in a microplate reader Infinite M1000 (Tecan Group AG). For testing of persistently HEV-infected cells, final ML2-SA1 concentrations ranging from 1 µM to 625 µM were used.

### 2.4 Time-of-addition experiment

A549 cells were seeded in six-well plates and infected with the ZIKV French Polynesia or Uganda isolate (MOI=1) for one hour, then unbound virus was removed. Treatment with final concentrations of 50 nM Bafilomycin A1, 100 µM ML-SA1, 100 µM ML2-SA1 or 100 µM Ribavirin was started at the respective time point (2 h before infection, during infection, 1 h, 6 h, 16 h and 22 h post infection). Complete media changes were performed during infection and 1 h post infection and compounds were refreshed, when applicable. At 24 h post infection, cells were washed once with PBS and lysed with RIPA (radio-immunoprecipitation assay) buffer for subsequent Western blot analyses.

### 2.5 Western blotting

A549 cells were washed with PBS, lysed with 100 µL RIPA buffer per well (including 10 µg/mL Aprotinin, 1 mM phenylmethylsulfonyl fluoride (PMSF), 25 µg/mL Leupeptin, 20 µg/mL Pepstatin (all purchased from AppliChem GmbH)) and scraped from plates. Afterwards, lysates were sonified and the protein concentration was determined using Bradford Reagent (Thermo Fisher Scientific). Samples containing 75 µg protein in SDS loading buffer were loaded on an SDS-PAGE (10% SDS) and separated using a voltage of 100 V. Afterwards, protein was transferred to a nitro-cellulose membrane by semi-dry western blotting. The membranes were blocked with 1X Roti-block (Carl Roth) for 1 h and incubated overnight with primary antibodies: anti-ZIKV E protein (Dr. Sami Akhras, Paul-Ehrlich-Institute (Akhras et al., 2019)) at a 1:500 dilution, anti-p62 (ProGen Biotechnik, 1:1,000), anti-LC3 (BIOZOL, 1:1,000), anti-pORF2 (HCD3K129, 1:2,000, raised against aa112-608 of recombinant pORF2 protein as used in a previous study (Glitscher et al., 2021a)) as well as anti-GAPDH (Santa Cruz Biotechnology, 1:2,000) and anti-beta-Actin (Sigma-Aldrich, 1:10,000 dilution) as loading control for quantification. The following day, membranes were washed with TBS (tris buffered saline) buffer containing 0.05% Tween and then incubated at a 1:10,000 dilution with secondary antibodies (IRDye® 680RD Donkey anti-Rabbit IgG, IRDye® 800CW Donkey anti-Guinea Pig IgG, IRDye® 800CW Donkey anti-Mouse IgG) purchased from LI-COR Biosciences.

### 2.6 RNA isolation, reverse transcription and RT-qPCR

A549 cells were washed with PBS lysed with RNA-Solv® Reagent (Carl Roth) and total intracellular RNA was isolated according to the manufacturer’s instructions. Afterwards, DNA was digested using the RQ1 RNAse-free DNase (Promega) and total RNA was transcribed into cDNA with random hexamer primer and RevertAid H Minus Reverse Transcriptase (Thermo Fisher Scientific) following the manufacturer’s protocol. ZIKV intracellular transcripts were quantified using the Maxima SYBR Green qPCR Kit (Thermo Fisher Scientific) for real-time quantitative PCR (RT-qPCR) in a LightCycler®480 System (Roche). The following primers were used for determination of intracellular ZIKV genomes (fwd 5’ agatcccggctgaaacactg 3’, rev 5’ ttgcaaggtccatctgtccc 3’). The housekeeping gene human ribosomal protein L27 (hRPL27, fwd 5’ aaagctgtcatcgtgaagaac 3’, rev 5’ gctgtcactttgcgggggtag 3’) was used for normalization.

### 2.7 Plaque assay

After the treatment of A549 cells with 100 µM ML2-SA1 and infection with ZIKV French Polynesia or ZIKV Uganda, supernatants were collected at the respective harvesting time point and stored at −80 °C. For plaque assays, Vero cells were seeded on a six-well plate (3x 10^5^ cells per well) and infected with serial dilutions of sample supernatant which was centrifuged at 5 min at 5000 rpm prior to remove cell debris. The inoculum was removed two hours post infection and cells were covered with 0.4% SeaPlaque^TM^ agarose (Lonza Group AG) in medium. After 20 min at room temperature (rt) for solidification of the agarose/medium mixture, cells were incubated in 5% CO_2_ and 95% relative humidity at 37 °C for five days. Plaque visualization was performed as described previously (Elgner et al., 2018).Virus titers are described as plaque forming units per mL (PFU/mL) and are presented in relation to the respective DMSO control.

### 2.8 Half maximal tissue culture infective dose (TCID_50_)

Determination of the TCID_50_ was performed as described previously (Glitscher et al., 2021a). In brief, highly HEV-permissive A549/D3 cells (Schemmerer et al., 2016) were infected using a serial dilution of HEV-containing cell culture supernatants replicates for 96 hours. Fixation and blocking were followed by incubation with an anti-pORF2 (HCD3K129) antibody, which was probed with a horseradish peroxidase-coupled secondary antibody (donkey-α-rabbit IgG, GE HealthCare, 1:400). Subsequent staining was performed using 3-amino-9-ethylcarbazol. The resulting TCID_50_ was calculated as described previously (Ramakrishnan, 2016).

### 2.9 Confocal laser scanning microscopy (cLSM) and super-resolution microscopy

Cells were seeded on coverslips (1.5H) (Carl Roth) and at the desired time-point fixed with ice-cold ethanol:acetone (1:1 v/v) for 10 min or 4% para-formaldehyde in PBS for 20 min at rt, respectively. For staining of acidic cellular organelles, 75 nM LysoTracker^TM^ Deep Red (Thermo Fisher Scientific) was added one hour before fixation. Blocking and permeabilization were performed by addition of 5% BSA in TBS-T for 15 min at rt. Samples were incubated with the following primary antibodies (anti-CD63 (Abcam, 1:200), anti-LAMP2 (R&D systems, 1:100), anti-LAMP2 (BD Biosciences, 1:200), anti-E (PEI, 1:500), anti-flavivirus group antigen antibody (clone D1-4G2-4-15, Sigma-Aldrich, 1:300), anti-dsRNA (SCICONS, 1:200), anti-pORF2 (HCD3K129, 1:500)) followed by an incubation step with the respective secondary antibodies (anti-mouse Cy^TM^3 (1:400), anti-rabbit IgG-Alexa 488 (1:1,000), anti-mouse IgG-Alexa 546 (1:1,000), anti-donkey IgG-Alexa 633 (1:1,000)) and DAPI (Carl Roth) for cell nuclei visualization. For staining of intracellular cholesterol and oxysterols, 0.1 mg/mL Filipin III (Sigma-Aldrich) in DMSO was added for a 30 min blocking step and incubation with secondary antibodies, respectively. Coverslips were mounted with mowiol on microscope slides and analyzed using a confocal Leica SP8 microscope (Leica). For super-resolution microscopy, samples were incubated with secondary antibodies (STAR635P (Abberior, 1:200) and Atto594 (Merck KgaA, 1:200)), which are suitable for STED (stimulated emission depletion). Image acquisition was performed on a Leica Stellaris 8 microscope system (Leica) equipped with a white light laser for fluorescence excitation and three STED lasers for fluorescence inhibition using a 93x objective (HC PL APO 93x/ 1.3 Glyc STED WHITE). For dual-color STED, a single STED line (775 nm) was used. The pinhole was set to 1.0 AU (airy unit). For z-stacks, step sizes were set to 0.2 µm. Immunofluorescence images were deconvoluted using the LasX Lightning tool. Acquisition and analysis of all images were performed using the LAS X software or FIJI. For determination of the thresholded Mender’s overlap coefficient (tMOC) or corrected total cell fluorescence (CTCF), a minimum of eight cells were analyzed.

### 2.10 ZIKV Luciferase Reporter Assay

A549 cells were seeded with a density of 2×10^5^ cells/well in a 12-well plate. The following day, cells were infected with 10 genome equivalents (GEs) of ZIKV Luciferase reporter virus for 4 h in the absence (DMSO only) or presence of 100 µM ML2-SA1/ML-SA1. At 4 h and 24 h post infection, compounds were refreshed to ensure a constant level throughout the experiment. At 48 h post infection, cells were washed, lysed with 130 µL of lysis buffer (PJK GmbH) and the Luciferase activity was determined using the Gaussia GLOW-Juice Kit (PJK GmbH) according to the manufacturer’s protocol. Luciferase activity was referred to the protein concentration of the corresponding lysate which was determined by a Bradford assay beforehand.

### 2.11 Statistical analyses

Results are depicted as mean of at least three independent experiments ± standard deviation (SD), unless stated otherwise. All data were analyzed using the software GraphPad Prism (Dotmatics) and student t test was used to analyze statistical relevance. The level of statistical relevance shown in the graphs are expressed with asterisks (*) that correspond to the following *p*-values: \**p* < 0.05, \*\**p* < 0.01, \*\*\**p* < 0.001, \*\*\*\**p* < 0.0001.

## 3 Results

### 3.1 The specific TRPML2 agonist ML2-SA1 displays an antiviral effect against different ZIKV isolates

A link between TRPML2 and flaviviral infections has been suggested previously (Rinkenberger and Schoggins, 2018), but the molecular details remained elusive. Based on the predominant localization of the cation channel in the endolysosomal system, we speculated that TRPML2 may play a role in the escape, maturation, or replication of endocytosed viruses and thus present a promising antiviral target (Fig. 1a).

**Figure 1:**
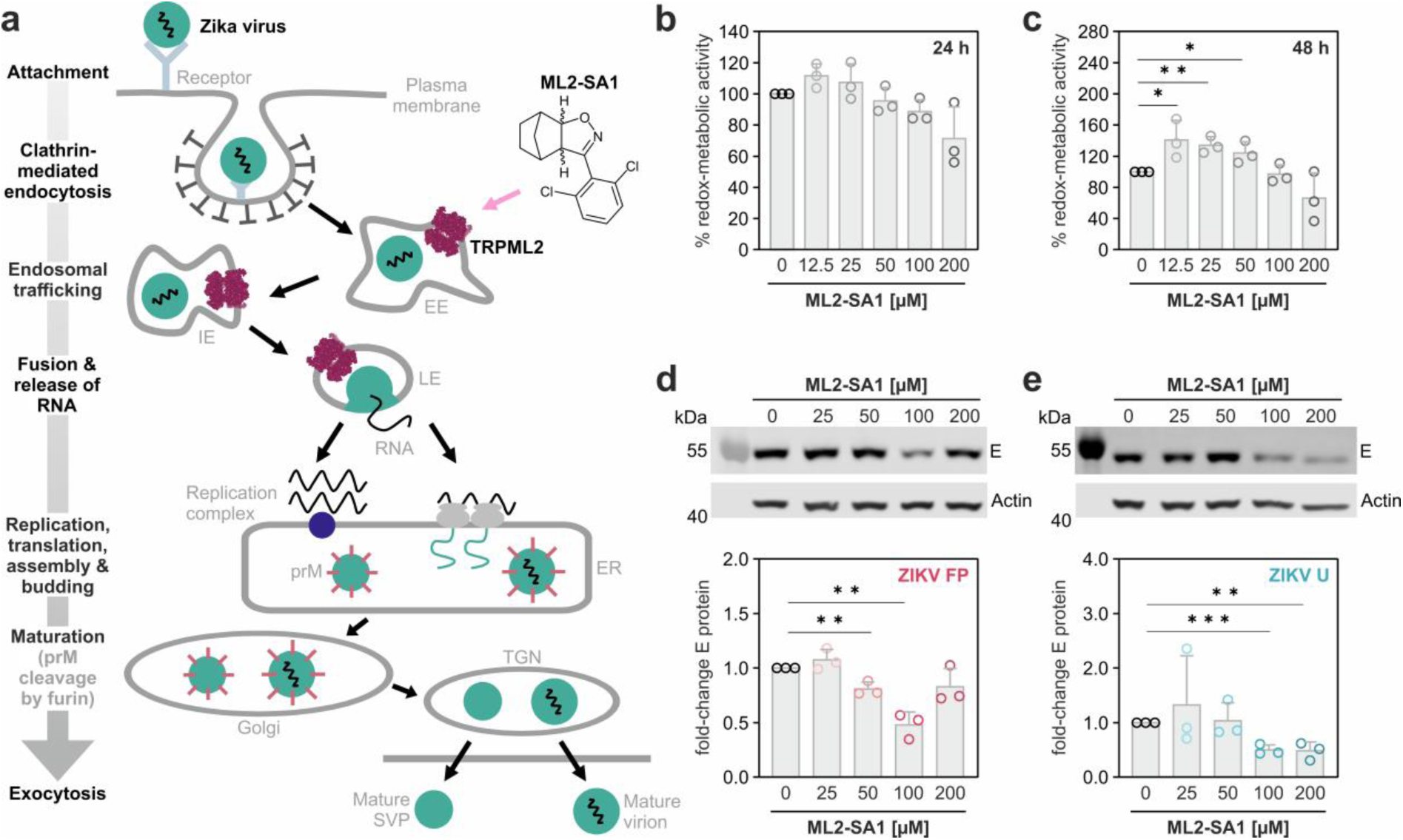
The specific TRPML2 agonist ML2-SA1 inhibits infection with two ZIKV isolates in a dose-dependent manner. **(a)** Schematic overview of the Zika virus life cycle. **Abbreviations:** EE: early endosome, IE: intermediate endosome, LE: late endosome, ER: endoplasmic reticulum, prM: pre-membrane protein, TGN: trans-Golgi network, SVP: subviral particle. **(b, c)** Cell viability of A549 cells upon treatment with 12.5 µM to 200 µM ML2-SA1 after 24 h (b) and 48 h (c) assessed with the PrestoBlue® assay. Values are expressed as % of intact cells normalized to the experimental control. **(d, e)** Relative fold-changes in ZIKV E protein of infected A549 cells treated with varying concentrations of ML2-SA1. ML2-SA1 was added to A549 cells two hours before infection (MOI = 1 with ZIKV French Polynesia (FP, H/PF/2013) or ZIKV Uganda (U, 976 Uganda)) and afterwards the compound was refreshed during every medium change to assure its constant presence. Cells were harvested 24 hpi followed by Western blotting to detect viral E protein. Data are normalized to the respective DMSO control and are expressed as mean ± SD from *n* = 3 biological replicates, a representative blot is shown. Statistical significance was determined by using an unpaired *t*-test. \**p* < 0.05, \*\**p* < 0.01, \*\*\**p* < 0.001, \*\*\*\**p* < 0.0001.

To investigate TRPML2’s role in ZIKV infections, we synthesized the only currently available specific TRPML2 agonist, ML2-SA1 (Plesch et al., 2018), according to published protocols ((McIntosh et al., 2012; Plesch et al., 2018), Scheme 1, Fig. S1–S4). Upon treatment of uninfected A549 cells with up to 100 µM ML2-S1, no significant cell toxicity was observed after 24 h (Fig 1b). Addition of 200 µM ML2-SA1 led to a mild but insignificant decrease in the redox metabolic activity after 24 and 48 h (Fig. 1b, c). After 48 h, an increase in the percentage of redox metabolic activity was detected upon addition of 12.5, 25 and 50 µM ML2-SA1 (Fig. 1c).

Next, we tested the antiviral potential of ML2-SA1 against ZIKV French Polynesia (H/PF/2013) and ZIKV Uganda (976 Uganda). At concentrations that do not induce toxicity in A549 cells, ML2-SA1 exhibited a dose-dependent reduction of the viral life cycle for both ZIKV isolates as gauged by the reduction in viral E protein (Fig. 1d, e). The most pronounced effect against both isolates was observed at a concentration of 100 µM ML2-SA1, which was thus used for all subsequent experiments.

To assess whether the reduction in the intracellular amount of E protein in ML2-SA1 treated ZIKV infected cells correlates with fewer intracellular genomes, quantification of the intracellular genomes and of the released viral particles was performed. For this purpose, ML2-SA1-treated A549 cells were infected with ZIKV French Polynesia or Uganda and harvested 24 h post infection (hpi) and 48 hpi, respectively. For both isolates, ML2-SA1 strongly reduced the amount of intracellular ZIKV E protein (Fig. 2a, b), as well as intracellular ZIKV genome levels (Fig. 2c, d). In accordance with this, the amount of released viral particles was also significantly diminished (Fig. 2e, f).

**Figure 2:**
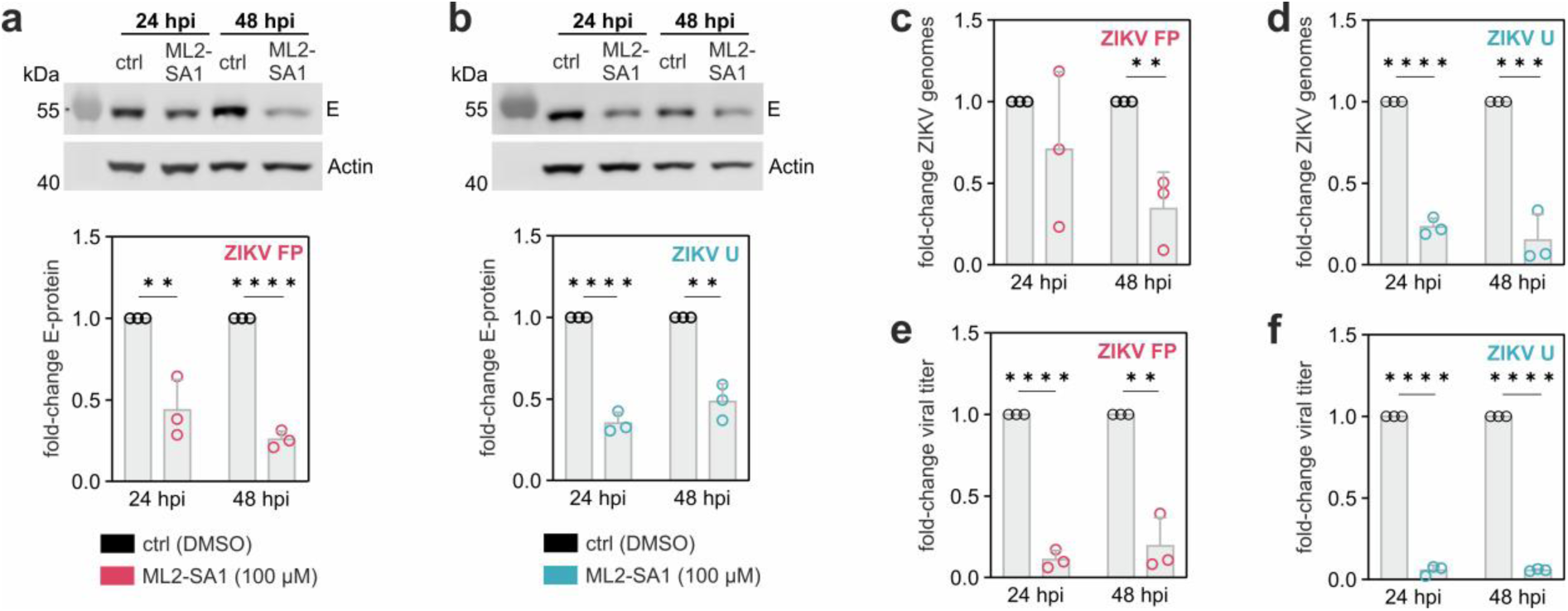
ML2-SA1 reduces intracellular ZIKV E protein and genome levels as well as the amount of released viral particles. A549 cells were treated with 100 µM ML2-SA1 for 2 h, then infected with the ZIKV French Polynesia (H/PF/2013) or ZIKV Uganda (976 Uganda) isolate (MOI = 1) and harvested 24 hpi and 48 hpi. (**a**, **b**) Relative fold-change of viral E protein determined by Western Blot. A representative Western Blot is shown above. (**c**, **d**) Relative fold-change in intracellular ZIKV genomes assessed via RT-qPCR. (**e**, **f**) Amount of released infectious ZIKV particles is determined using a plaque assay. Data are normalized to the respective DMSO control for each time point and are expressed as mean ± SD from *n* = 3 biological replicates. Statistical significance was determined by using an unpaired *t*-test. \**p* < 0.05, \*\**p*< 0.01, \*\*\**p* < 0.001, \*\*\*\**p* < 0.0001.

Taken together, these data indicate that the TRMPL2 agonist ML2-SA1 exerts a significant antiviral effect on ZIKV.

### 3.2 ML2-SA1 treatment causes accumulation of ZIKV E protein in CD63-positive vesicles without leading to enhanced ZIKV degradation

With its ability to reduce ZIKV genome levels and virus titers, the TRPML2 agonist displayed an appreciable antiviral effect against both tested ZIKV isolates. However, it remained unclear whether the compound interferes with early infection stages (e.g. through increased lysosomal degradation of ZIKV thereby decreasing viral RNA release into the cytosol) or later infection stages (i.e. impairment of viral replication). To investigate whether ML2-SA1 treatment influences the pH of endolysosomal vesicles, uninfected A549 cells were subjected to ML2-SA1 or the V-type ATPase inhibitor Bafilomycin A1, which prevents endolysosomal acidification (Fig S5). Acidic organelles were stained with the fluorescent dye LysoTracker^TM^ and immunostaining of the lysosomal marker LAMP2 was carried out. As expected, almost no LysoTracker^TM^ positive vesicles were found in Bafilomycin A1-treated cells. In contrast, ML2-SA1-treated cells displayed more and enlarged LysoTracker^TM^-positive vesicles compared to the DMSO-treated control. In addition, a drastic increase in the number as well as size of LAMP2-positive vesicles was observed for ML2-SA1-treated cells. These results indicate that treatment with the TRPML2 agonist ML2-SA1 influences either the morphology of or fusion events within the endolysosomal system.

To investigate whether ML2-SA1 treatment induces accumulation of ZIKV particles in the endolysosomal system, a prerequisite for a possible increase in lysosomal ZIKV degradation, we performed super-resolution microscopy (Fig. 3a). Infected cells were immunostained against ZIKV E protein and human CD63, a marker for multivesicular bodies (MVBs). In untreated cells, the ZIKV E protein signal was concentrated around the nucleus, in agreement with the formation of ER-derived structures associated with virus replication and particle assembly (Cortese et al., 2017). In ML2-SA1-treated cells, the ZIKV E protein signal was reduced and more evenly distributed across the cell suggesting that replication was reduced or halted. Furthermore, ML2-SA1 treatment increased the size of CD63-positive MVBs in comparison to the DMSO control. Representative 3D reconstructions revealed that ZIKV E protein accumulated in these CD63-positive vesicles upon ML2-SA1 treatment.

**Figure 3:**
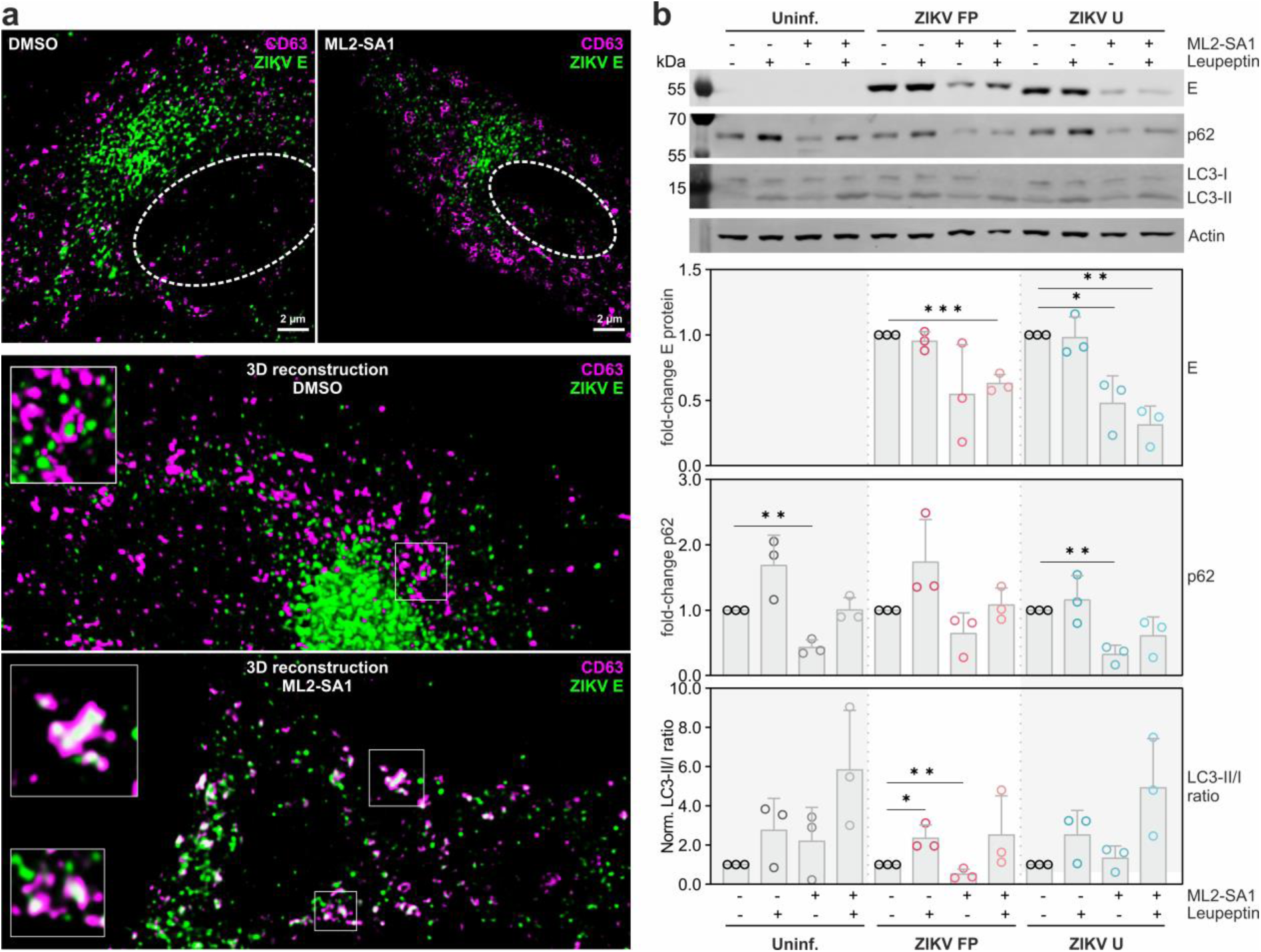
TRPML2 agonist treatment leads to ZIKV E protein accumulation in multivesicular bodies but not increased lysosomal degradation. **(a)** Representative super resolution microscopy images of A549 cells treated with DMSO (left) or 100 µM ML2-SA1 in DMSO (right) and infected with ZIKV Uganda (976 Uganda). Cells were fixed 24 hpi for immunostaining with antibodies against ZIKV E protein (green) and CD63 (magenta). The position of the cell nucleus is indicated by a dotted circle. Scale bar: 2 µm. Representative 3D reconstructions are shown below. **(b)** Relative fold-change of E protein, p62 and LC3 determined by Western Blotting. Representative Western Blots are shown above. A549 cells were treated with 100 µM ML2-SA1 and/or 0.1 mg/mL Leupeptin for 2 h before infection. Cells were infected (MOI = 1) with the ZIKV French Polynesia (H/PF/2013) and ZIKV Uganda (976 Uganda) isolate and harvested 24 hpi and 48 hpi. Data are normalized to the respective DMSO control and are expressed as mean ± SD from *n* = 3 biological replicates. Statistical significance was determined by using an unpaired *t*-test. \**p* < 0.05, \*\**p* < 0.01, \*\*\**p* < 0.001, \*\*\*\**p* < 0.0001.

A prior study suggested increased lysosomal degradation of DENV2 and ZIKV upon treatment of infected cells with the unspecific TRPML-agonist ML-SA1 (Xia et al., 2020). Therefore, we investigated whether ML2-SA1-treatment increases ZIKV lysosomal degradation. We incubated infected A549 cells with either ML2-SA1, the membrane-permeable thiolprotease inhibitor Leupeptin, or both compounds. Leupeptin inhibits the lysosomal proteases Cathepsin B, H and L thereby decreasing the proteolytic degradation of lysosomal cargo (Furuno et al., 1982; Glaumann and Ahlberg, 1987). While Leupeptin alone had no effect on ZIKV E protein levels (Fig. 3b), the co-treatment of ML2-SA1 and Leupeptin also showed no increase of intracellular ZIKV E protein.

As a control for the efficiency of lysosomal proteolysis, we thus checked whether Leupeptin application resulted in the expected decrease in degradation of another lysosomal marker protein, p62 (Fig. 3b). The observed increase in p62 levels upon Leupeptin addition confirms that inhibition of the acidic lysosomal proteases indeed leads to reduced lysosomal protein degradation. Interestingly, the combination of Leupeptin with ML2-SA1 showed no statistically significant changes in p62 levels, however when cells were treated with ML2-SA1 alone, a statistically significant p62 decrease was observed in both uninfected and ZIKV Uganda-infected cells (Fig. 3b) which might reflect the enhanced lysosomal capacity. Since a decrease in p62 levels has also been associated with enhanced autophagy (Bjørkøy et al., 2009), we additionally looked at the increase of the LC3-II/I ratio, a second autophagy marker, which would imply enhanced autophagosome formation (Kabeya et al., 2004). Here, only the treatment of ZIKV French Polynesia infected cells with ML2-SA1 showed a significant effect (Fig. 3b). Together, these data suggest that lysosomal degradation is not the underlying molecular mechanism of the observed antiviral effect of ML2-SA1 and that enhanced autophagy plays only a minor role.

### 3.3 Viral replication, and not the early ZIKV life cycle stages, are impaired by ML2-SA1 treatment

Time-of-addition experiments allow the dissection which step of the viral life cycle is affected by a given compound (Daelemans et al., 2011) (Fig 4a). For instance, Bafilomycin A1 restricts ZIKV entry by inhibiting the V-type ATPase (Cortese et al., 2017; Sabino et al., 2019). Ribavirin, a nucleoside analogue, inhibits ZIKV replication (Kamiyama et al., 2017) and the non-specific TRPML channel agonist ML-SA1 was described to enhance lysosomal degradation of ZIKV (Xia et al., 2020).

**Figure 4:**
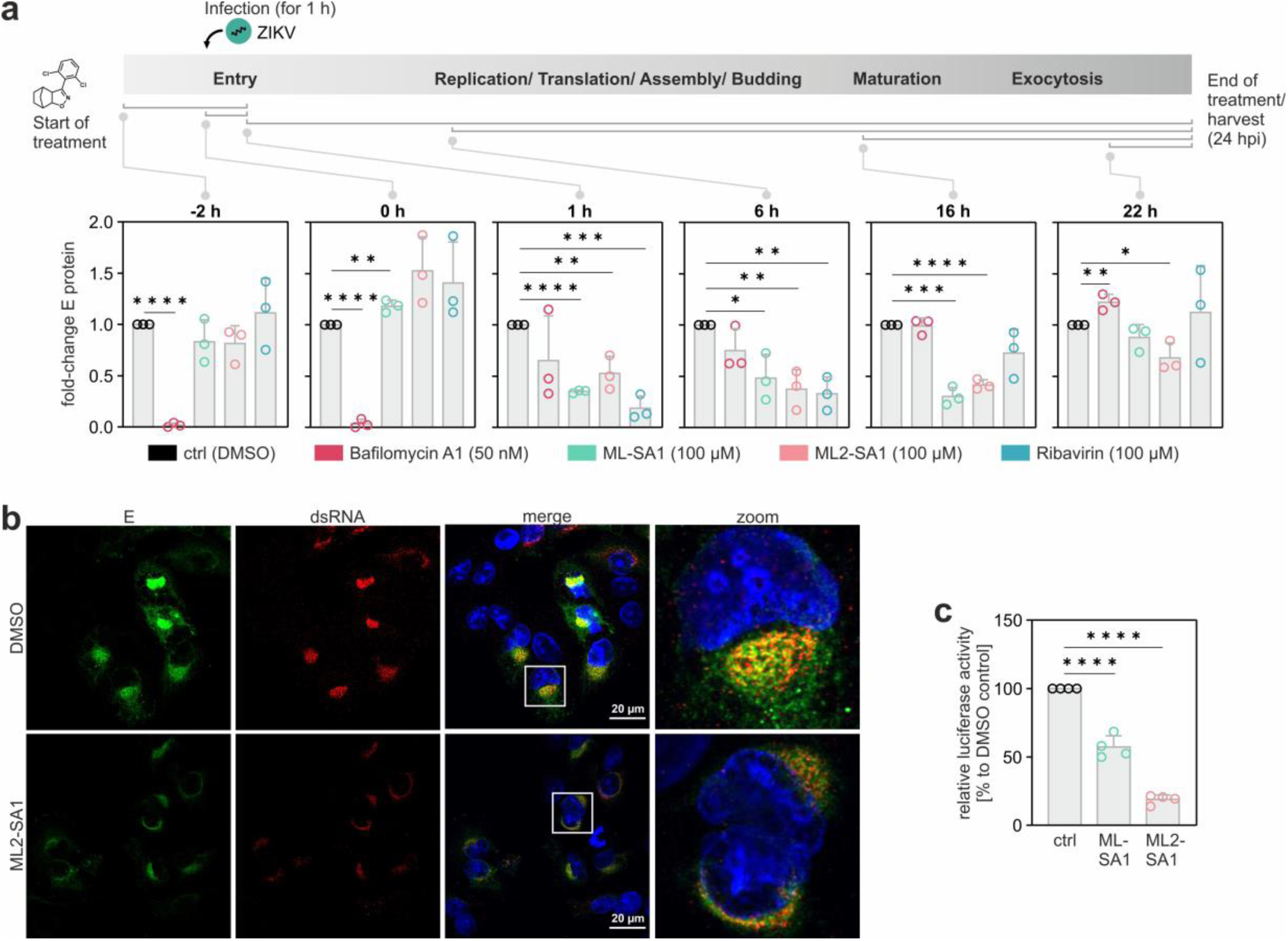
TRPML2 agonist treatment affects late stages of the ZIKV life cycle. **(a)** A549 cells were infected with ZIKV Uganda isolate (976 Uganda) for one hour (MOI = 1), then unbound virus was removed. Treatment with either 50 nM Bafilomycin A1, 100 µM ML-SA1, 100 µM ML2-SA1 or 100 µM Ribavirin was started at the respective time points, i.e. −2h before infection, during infection, as well as 1 h, 6 h, 16 h or 22 h after infection. Gray brackets below the time scale indicate the respective compound incubation period. The resulting relative fold-change of ZIKV E protein was assessed by Western Blotting. Representative Western Blots are shown in Fig. S6a. Results are normalized to the respective DMSO control of each time point and are expressed as mean ± SD from *n* = 3 biological replicates. **(b)** Representative confocal laser scanning microscopy (cLSM) images of A549 cells treated with 100 µM ML2-SA1 for 2 h before infection. Cells were infected (MOI = 1) with the ZIKV Uganda (976 Uganda) isolate and fixed 24 hpi. Nuclei were stained with DAPI (blue). ZIKV E protein (green) and ZIKV dsRNA (red) were visualized using specific antibodies. Scale bar: 20 µm. **(c)** A549 cells were treated with 100 µM ML-SA1 or 100 µM ML2-SA1 and simultaneously infected with ZIKV *R*Luc reporter virus for 4 h. Compounds were refreshed during each media change. 48 h post infection, Luciferase activity was determined. Results are normalized to the DMSO control and are expressed as mean ± SD from *n* = 4 biological replicates. Statistical significance was determined by using an unpaired *t*-test. \**p* < 0.05, \*\**p* < 0.01, \*\*\**p* < 0.001, \*\*\*\**p* < 0.0001.

To dissect the consequence of ML2-SA1 treatment, A549 cells were infected with the ZIKV Uganda isolate or ZIKV French Polynesia and the TRPML2 agonist was added at defined time points (from −2 to +22 hpi) to capture the entry and post entry phase of the viral life cycle (Fig. 4a, Fig. S6). To benchmark the effects of the TRPML2 channel agonist to specific events of the ZIKV life cycle stage, cells were also treated with Bafilomycin A1, Ribavirin and ML-SA1. Bafilomycin A1 and Ribavirin were used at concentrations known to be non- or only mildly cytotoxic, i.e. 50 nM and 100 µM, respectively (Huber et al., 2022; Sabino et al., 2019). When testing the cytotoxicity of ML-SA1 in the same manner as we did for ML2-SA1, we found that neither compound displayed significant cytotoxicity at a concentration of 100 µM (Fig. S7), the concentration consequently used in our time-of-addition assay for both molecules.

In the assay, the amount of detected ZIKV E protein was normalized to the respective DMSO control (Fig. 4a, black circles) to monitor the relative changes in the amount of viral E protein for each compound. In agreement with the literature, we found Bafilomycin A1 to strongly suppress viral entry as reflected by the significant reduction in viral E protein at early addition time points (Fig. 4a, magenta circles). Ribavirin affected the later stages of the viral life cycle, 1–6 hpi, including viral replication (Fig. 4a, blue circles). Both ML2-SA1 and ML-SA1 acted similar to Ribavirin and showed no antiviral effect when added during the entry process, but at later stages, (Fig. 4a, green and light pink circles). No apparent differences between the experiments with ZIKV Uganda or ZIKV French Polynesia isolate were observed (see Fig. S6b, c for the time-of-addition experiment with ZIKV French Polynesia infected cells).

Since we found that ML2-SA1 did not increase the amount of released viral particles (Fig. 2e, f) and not lead to enhanced ZIKV protein degradation in the endolysosomal system (Fig. 3), it seemed most likely that the observed antiviral activity in the time-of-addition experiment originates from an impairment in ZIKV replication. The key ZIKV replication stages (translation and cleavage of the viral polyprotein, replication of viral RNA and formation of progeny ZIKV particles) take place at the ER of the host cell (Cortese et al., 2017; Rossignol et al., 2017). During the formation of these viral ‘replication factories’, both the ER and cytoskeleton undergo massive remodeling (Cortese et al., 2017; Long et al., 2020; Rossignol et al., 2017). Dense, ER-derived patches in the perinuclear region which give rise to dsRNA intermediates during ZIKV replication can be observed (Cortese et al., 2017). In addition, these cellular changes result in a characteristic bean-like shape formation of the host cell nucleus (Cortese et al., 2017). Using confocal laser scanning microscopy, we observed the characteristic re-shaping of host cell nuclei and large, dsRNA-positive patches in the perinuclear region in ZIKV-infected A549 cells (Fig. 4b). ML2-SA1 treatment caused a significant decrease of dsRNA in ZIKV Uganda isolate infected A549 cells. Furthermore, the remaining dsRNA signal was not concentrated in distinct perinuclear patches anymore, but rather evenly distributed. This suggests that the TRPML2 agonist prevented the formation of pronounced ZIKV replication sites. As described above, ML2-SA1 also caused a decrease in the number of ZIKV genomes (Fig. 2c, d). The notion that ML2-SA1 impairs ZIKV replication was also supported by a ZIKV *Renilla* Luciferase (*R*Luc) reporter assay, which takes advantage of a *R*Luc gene fused to the viral capsid coding sequence (Shan et al., 2016). Here, treatment of A549 cells with ML-SA1 or ML2-SA1 resulted in a significant decrease of Luciferase activity for both compounds. Interestingly, this effect was more pronounced for ML2-SA1 compared to ML-SA1 (Fig. 4c).

### 3.4 ML2-SA1 treatment leads to intracellular cholesterol accumulation

Although a functional role for TRPML1 in the regulation of lysosomal lipid and cholesterol trafficking has been described (Shen et al., 2012), and the cryo-electron microscopy structure of TRPML3 displays bound cholesterol molecules (Hirschi et al., 2017), no obvious connection between TRPML2 and cholesterol binding or trafficking has been established to date. Since efficient ZIKV replication requires cholesterol, which affects membrane rigidity and induces negative membrane curvature, to enable the formation or maintenance of flaviviral replication factories (Churchward et al., 2005; Steinkühler et al., 2019), we sought to explore whether addition of TRPML channel agonists may also affect intracellular cholesterol levels (Fig. 5). Using immunofluorescence, we did not observe distinct differences of cellular cholesterol distribution between infected and uninfected cells. However, upon treatment with ML2-SA1, regardless of whether cells were infected or not, pronounced cholesterol accumulation was observed in dot-like structures. In contrast, cholesterol was observed as a dispersed cytosolic signal with small punctae in untreated cells. Consistent with our previous findings (Fig. S5), ML2-SA1-treatment also enlarged the size of LAMP2-positive structures. Notably, treatment with the nonspecific TRPML channel agonist ML-SA1 had a similar effect, both with regards to cholesterol redistribution and enlarged vesicular structures. It remains to be seen whether this can solely be attributed to the molecule’s action on TRPML2, or also on other TRPML channels. In summary, our results indicate that both ML-SA1 and ML2-SA1 alter the intracellular cholesterol distribution in a similar manner which may play a role in the observed antiviral effects.

**Figure 5:**
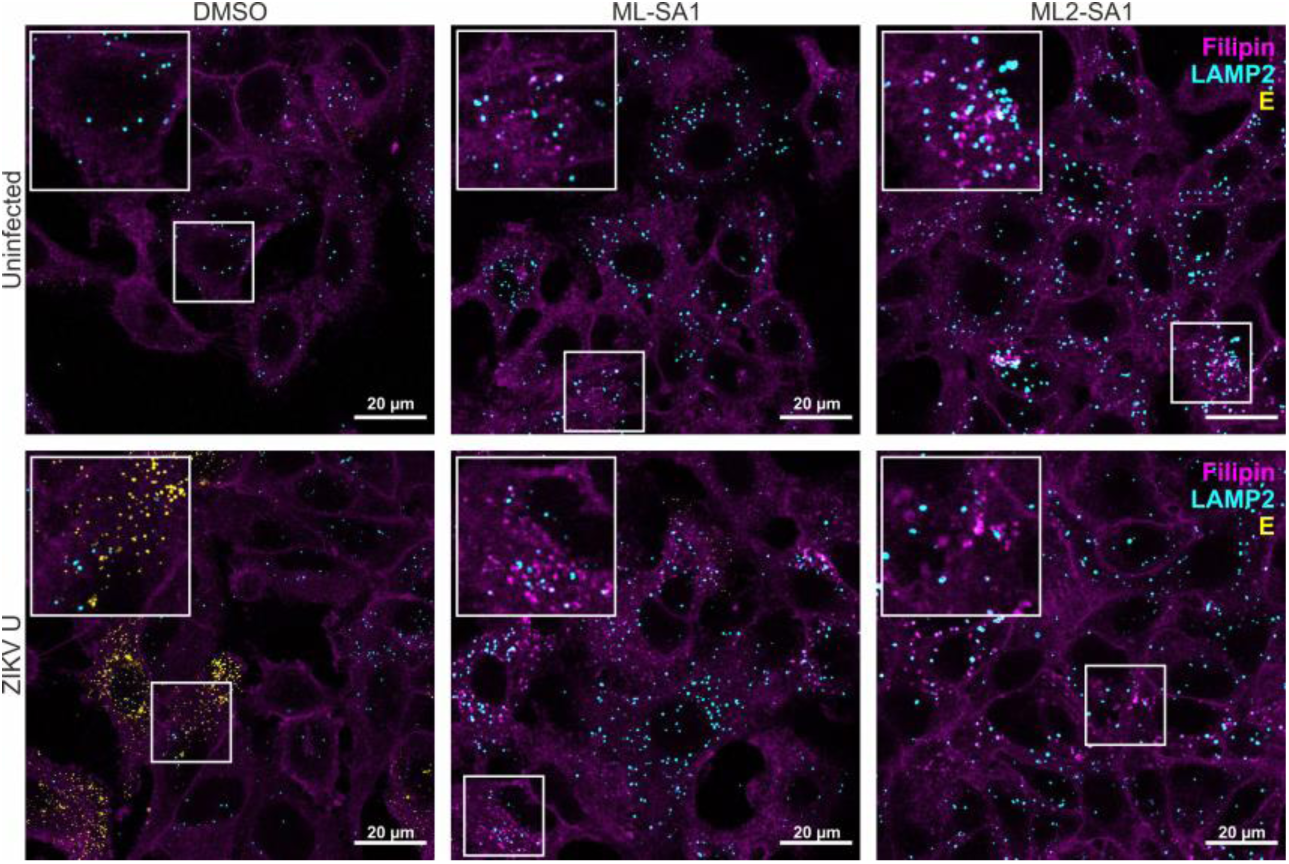
Treatment with ML-SA1 or ML2-SA1 leads to cholesterol accumulation in uninfected and infected cells. Representative confocal laser scanning microscopy images of A549 cells treated with 100 µM ML-SA1 or 100 µM ML2-SA1 for 2 h before infection. Cells were infected (MOI = 1) with the ZIKV Uganda (976 Uganda) isolate and fixed 24 hpi. Intracellular cholesterol and oxysterols were stained using Filipin (magenta). ZIKV E protein (yellow) and the lysosomal marker LAMP2 (cyan) were visualized using specific antibodies. Scale bar: 20 µm.

### 3.5 ML2-SA1 enhances lysosomal degradation and decreases release of functional HEV virions

TRP channels in general and lyososmal Ca^2+^ channels specifically have been associated with viral infections beyond the flavivirus genus (Grimm et al., 2017; Kumar et al., 2022). We thus sought to investigate whether the TRPML2 agonist ML2-SA1 is also effective against endocytosed viruses beyond ZIKV. To this end, A549 cells persistently infected with HEV were treated with ML2-SA1. HEV is the most common cause of acute viral hepatitis and treatment options are still limited (Kamar et al., 2017). Interestingly, elevated intracellular cholesterol levels promote lysosomal degradation of HEV, while reduced intracellular cholesterol levels increase viral release (Glitscher et al., 2021b). In our hands, ML2-SA1 displayed only mild cytotoxicity in persistently HEV-infected A549 cells (Fig. 6a), but significantly decreased viral titers (Fig. 6b) alongside a heavy reduction in the amount of intracellular capsid protein pORF2 (Fig. 6c). This decrease in pORF2 was confirmed on a single-cell level by cLSM analyses (Fig. 6d, e). Most importantly, there are clear signs of the capsid protein being incorporated into lysosomal structures harboring LAMP2 (Fig. 6f). This suggests an increase in lysosomal degradation of HEV upon treatment with ML2-SA1, which likely is due to the cholesterol accumulation induced by ML2-SA1 treatment (Fig. 5).

**Figure 6:**
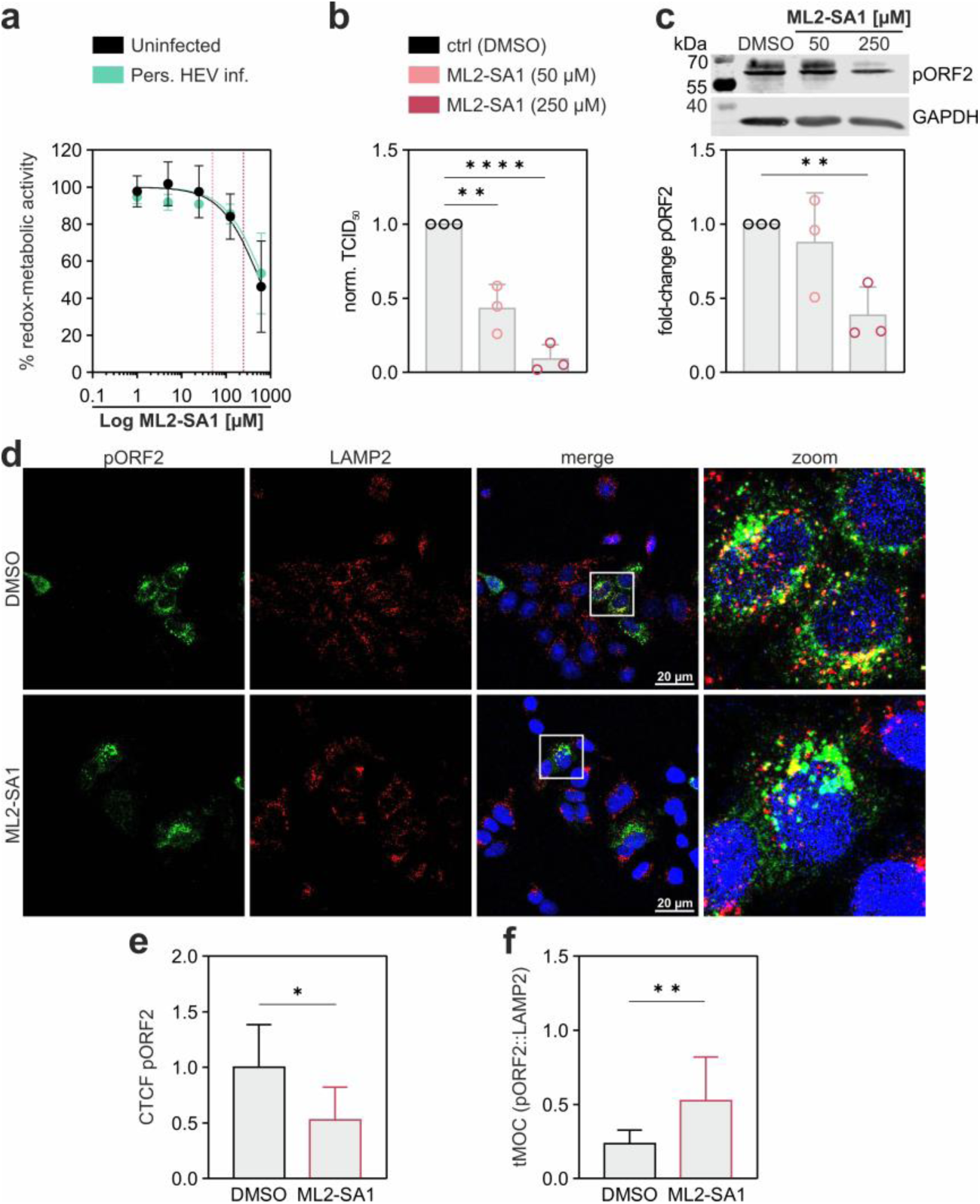
ML2-SA1 reduces intracellular HEV capsid protein by increasing lysosomal degradation and decreases the number of released virions. **(a)** Cell viability of uninfected (black) and persistently HEV-infected (green) A549 cells after treatment with 1 µM to 625 µM ML2-SA1 for 24 h assessed with the PrestoBlue® assay. Values are expressed as % of intact cells normalized to the experimental control. ML2-SA1-concentrations of 50 µM (light pink) and 250 µM (dark pink) are indicated by dotted lines. **(b)** Amount of released infectious HEV particles. **(c)** Relative fold-changes in HEV pORF2 protein of persistently infected A549 cells treated with different concentrations of ML2-SA1. Cells were harvested 24 h post-treatment followed by Western blotting to detect viral pORF2. Data are normalized to the respective DMSO control and are expressed as mean ± SD from *n* = 3 biological replicates, a representative blot is shown above. Statistical significance was determined by using an unpaired *t*-test. \**p* < 0.05, \*\**p* < 0.01, \*\*\**p* < 0.001, \*\*\*\**p* < 0.0001. **(d)** Representative confocal laser scanning microscopy (cLSM) images of persistently HEV-infected A549 cells treated with 250 µM ML2-SA1. Cells were fixed 24 h post treatment. Nuclei were stained with DAPI (blue), HEV pORF2 (green) and LAMP2 (red) were visualized using specific antibodies. Scale bar: 20 µm. **(e)** Normalized signal intensities of HEV pORF2 in micrographs depicted in (d) calculated by corrected total cell fluorescence (CTCF). Data are expressed as mean ± coefficient of variation (CV). **(f)** Absolute degree of co-localization between pORF2 and LAMP2 in (d) determined by calculation of thresholded Mender’s overlap coefficient (tMOC); 0 represents no colocalization; 1 represent a complete overlap of signals. Data are expressed as mean ± CV.

## 4 Discussion

Both TPCs (two-pore channels) and TRPML channels are endolysosomal cation channels which play important roles in viral infections (Chao et al., 2023; Grimm et al., 2017; Huang et al., 2021; Kumar et al., 2022; Rosato et al., 2021; Sakurai et al., 2015; Santoni et al., 2020). However, their antiviral potential is not yet fully explored, in part because specific functional modulators are missing. The recent synthesis and functional characterization of the first TRPML2-specific agonist, ML2-SA1, by Grimm and colleagues (Plesch et al., 2018) provided us with the unique opportunity to assess the role of this ion channel in the ZIKV infection process (Fig. 7).

**Figure 7:**
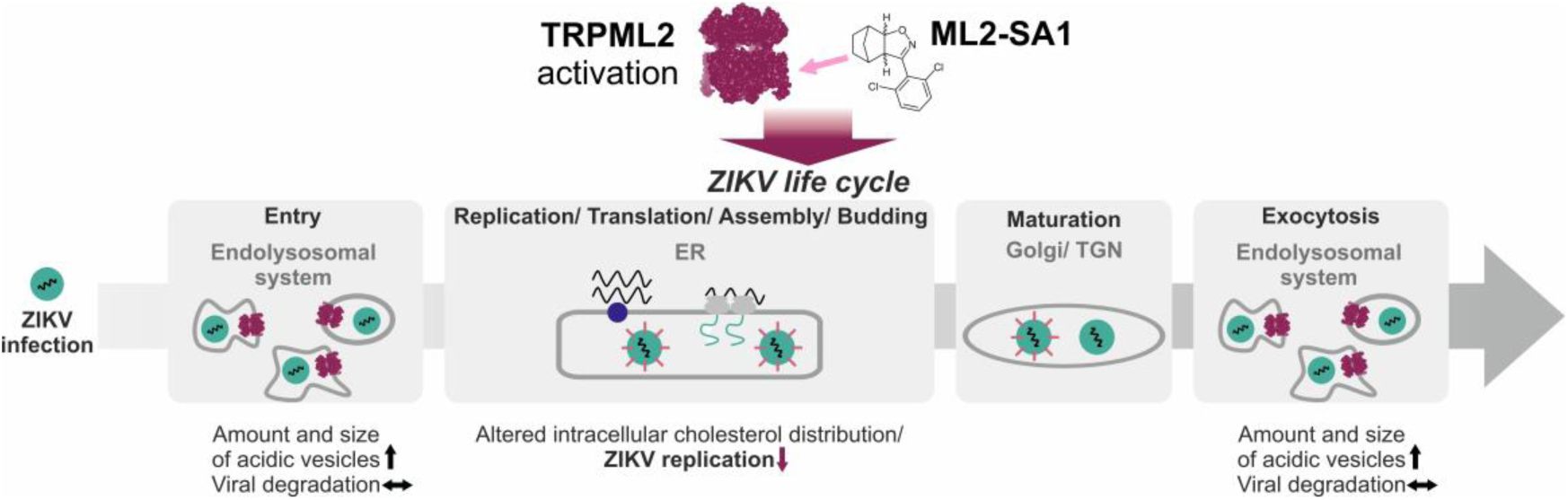
Specific TRPML2 agonist affects multiple stages of the ZIKV life cycle. Treatment with the TRPML2 agonist ML2-SA1 caused increased acidification of the endolysosomal system, however, enhanced lysosomal degradation of ZIKV was not the basis of the antiviral effect of TRPML2 activation. Instead, ML2-SA1 impaired ZIKV replication, presumably by an altered intracellular cholesterol distribution.

For the non-specific TRPML agonist ML-SA1, which shows antiviral potential against ZIKV and DENV2 in A549 cells, lysosomal acidification, increased lysosomal protease activity and enhanced viral lysosomal degradation were proposed to be the main mechanisms of action (Xia et al., 2020). However, the fact that ML-SA1 addresses TRPML channels indiscriminately and that its mechanism of action has not yet been investigated compared to reference compounds in time-of-addition experiments makes it impossible to disentangle the roles of agonist treatment and the activity of individual TRPML subfamily members in flaviviral infections. Using the specific TRPML2 agonist ML2-SA1, we encouragingly observed significant antiviral activity for ML2-SA1 in the same concentration range previously used for ML-SA1 against DENV and ZIKV (Xia et al., 2020).

We also observed an increase in amount and size of acidic vesicles and accumulation of ZIKV E protein in CD63-positive particles in ML2-SA1-treated cells in agreement with the prior study on ML-SA1. However, our synchronized time-of-addition experiments, the use of reference compounds and the inhibition of lysosomal proteases strongly suggested that the antiviral effect of this compound is not rooted in enhanced acidification and viral protein degradation (Fig. 3b, Fig. 4). Rather, in our hands both ML2-SA1 and ML-SA1 affected the late stages of the viral life cycle, similar to the replication inhibitor Ribavirin.

Interestingly, for both ML-SA1 and ML2-SA1, we also observed slightly proviral effects upon addition at early infection stages. Activation of TRPML channels leads to an increased Ca^2+^ flux into the cytosol which can enhance endolysosomal trafficking (Venkatachalam et al., 2015) and may therefore promote viral infection during the early stages of the ZIKV life cycle. We saw that ML2-SA1 addition resulted in MVBs with an increased size (Fig. 3a) which could indicate that TRPML2 activation increased vesicle trafficking and membrane fusion. This finding also agrees with the observations by Rinkenberger and Schoggins using a TRPML2 overexpression system, where enhanced endosomal trafficking led to a proviral effect for multiple endocytosed viruses including ZIKV (Rinkenberger and Schoggins, 2018).

We hypothesized that the pronounced impact of ML2-SA1 on ZIKV replication may be due to the prevention of the structural remodeling of the ER and impairment of the formation or maintenance of ZIKV replication factories, both processes that are contingent on cholesterol (Osuna-Ramos et al., 2018). Indeed, we found an altered intracellular cholesterol distribution upon treatment with ML2-SA1 (Fig. 4 and Fig. 5), thereby establishing the first link between TRPML2 activity and cellular cholesterol distribution. Ca^2+^ channel modulators have previously been shown to reduce ZIKV replication by interfering with intracellular cholesterol trafficking. Lacidipine for example inhibits L-type calcium channels located in the plasma membrane of vascular smooth muscle cells (Godfraind and Salomone, 1991; Spampinato et al., 1993) and interferes with intracellular trafficking of cholesterol and other lipids thereby inhibiting ZIKV replication (Bezemer et al., 2022). Host-cholesterol is also an important factor during viral entry (Glitscher and Hildt, 2021; Osuna-Ramos et al., 2018). The NPC1 inhibitor U18666A inhibits both DENV entry via retarded viral trafficking in cholesterol-loaded late endosomes/lysosomes and viral replication (Poh et al., 2012). During infection with the Hepatitis C virus (HCV), U18666A was shown to disturb the lipid transport, which causes functional inhibition of late endosomes and formation of multilamellar bodies (Elgner et al., 2016). In ZIKV-infected cells, U18666A treatment specifically inhibited viral replication (Sabino et al., 2019). It is therefore tempting to speculate that changes in cellular cholesterol distribution are also the molecular basis for the observed antiviral effect of ML2-SA1. Such a broad mechanism of action would imply that the antiviral potential of ML2-SA1 does not end with ZIKV but should extend to other viruses reliant on cholesterol. Indeed, many viruses depend on cholesterol during viral entry (Glitscher and Hildt, 2021; Osuna-Ramos et al., 2018) and elevation of intracellular cholesterol was successfully used to reduce infection with HEV by increasing lysosomal degradation of viral pORF2 (Glitscher et al., 2021b). Encouragingly, ML2-SA1 also showed antiviral activity against HEV (Fig. 6).

In this study we saw that the TRPML2-specific agonist ML2-SA1 induced a pronounced antiviral effect against two ZIKV isolates, one representative of the African lineage (ZIKV Uganda) and one representative of the Asian lineage (French Polynesia). ML2-SA1 treatment did not induce increased lysosomal ZIKV degradation, but rather inhibited viral replication, possibly due to an altered intracellular cholesterol distribution. Due to the high similarity of the life cycle of flaviviruses, it seems likely that ML2-SA1 also shows antiviral potential against other flaviviruses such as DENV, WNV and yellow fever virus (YFV). It will be interesting to investigate the connection between TRPML2, the other TRPML channels, viral infections, and host cell cholesterol distribution in more detail. It may be well conceivable that all members of the TRPML channel subfamily play central regulatory roles within the lipid distribution networks of the cell and that this functionality could provide a lever for the development of novel antiviral drugs targeting endolysosomal TRP ion channels.

## Author contributions

Experiment design, K.K.S, M.G., E.H., and U.A.H.; cell viability assays, western blotting, K.K.S., S.P., and M.G., time-of-addition experiment, RT-qPCR and plaque assay, K.K.S., TCID_50_, S.P. and M.G.; microscopy, K.K.S., D.B. and M.G.; ZIKV Luciferase reporter assay, R.M. and D.B.; organic synthesis, K.S.; data analysis and interpretation, K.K.S., M.G., D. B., E.H. and U.A.H.; writing of the paper, K.K.S., M.G., E.H. and U.A.H. with contributions from all authors; study supervision, T.S., E.H., and U.A.H.

## Acknowledgements

We thank Gert Carra for technical support, and Charlotte Guhl for fruitful discussions. K.K.S. acknowledges a PhD fellowship of the Max Planck Graduate Center (MPGC). U.A.H. acknowledges funding through a Boehringer Ingelheim Foundation Exploration Grant. This work was supported by the Deutsche Forschungsgemeinschaft (DFG, German Research Foundation) through the collaborative research center 1507 “Membrane-associated Protein Assemblies, Machineries, and Supercomplexes” (Project ID 450648163), the collaborative research center 1278 “Polymer-based nanoparticle libraries for targeted anti-inflammatory strategies” (project ID 316213987) and the Cluster of Excellence “Balance of the Microverse” (EXC 2051—Project-ID 390713860) (to U.A.H.). E.H. obtained a grant by the LOEWE Center DRUID (Novel Drug Targets against Poverty-Related and Neglected Tropical Infectious Diseases; Project D2).

## Abbreviations

aa: amino acids
cLSM: confocal laser scanning microscopy
CTCF: corrected total cell fluorescence
CV: coefficient of variation
DENV: Dengue virus
DMSO: dimethyl sulfoxide
E: protein envelope protein
FBS: fetal bovine serum
GEs: genome equivalents
HCV: Hepatitis C virus
HEV: Hepatitis E virus
hpi: hours post infection
IAV: Influenza A virus
ML2-SA1: mucolipin 2 synthetic agonist 1
ML-SA1: mucolipin synthetic agonist 1
MOI: multiplicity of infection
MVB: multivesicular body
PBS: phosphate buffered saline
PFU: plaque forming units
PIFA: (bis(trifluoroacetoxy)iodo)benzene
PMSF: phenylmethylsulfonyl fluoride
*Renilla*: Luciferase *R*Luc
RIPA: radio-immunoprecipitation assay
rt: room temperature
RT-qPCR: real-time quantitative polymerase chain reaction
SD: standard deviation
STED: stimulated emission depletion
TBS: tris buffered saline
TCID_50_: half maximal tissue culture infective dose
tMOC: thresholded Mender’s overlap coefficient
TPC: two-pore channels
TRPML: transient receptor potential mucolipin
WNV: West Nile virus
YFV: yellow fever virus
ZIKV: FP ZIKV French Polynesia isolate
ZIKV: U ZIKV Uganda isolate
ZIKV: Zika virus

