## Supplementary information for "Late stages of the Zika virus life cycle are impaired by a selective TRPML2 agonist"

### Chemistry

All solvents and reagents were obtained from commercial suppliers (Sigma-Aldrich, Acros Organics, Alfa Aesar, TCI, or Apollo Scientific) and used without prior purification. Reaction progress was monitored by thin-layer chromatography (TLC) using Machery-Nagel Alugram Xtra Sil G/$UV_{254}$ silica 60 plates. Compound visualization on these plates was attained by radiation at 254 nm. Purification by flash chromatography were performed on silica gel (0.015−0.040 mm, Machery-Nagel). $^{1}$H and $^{13}$C NMR spectra were recorded on a Bruker Fourier 300. All chemical shifts were referenced to the signal of the residual solvent ($CDCl_3$, 7.26 and 77.16 ppm; DMSO-$d_6$, 2.50 and 39.52 ppm for $^{1}$H NMR and $^{13}$C NMR, respectively) and reported in parts per million (ppm) relative to tetramethylsilane (TMS). Infrared (IR) spectra were recorded on an FT-IR spectrometer (Avatar, Thermo-Nicolet) with an ATR correction and are reported in terms of frequency of absorption $\bar{\nu}$ [$cm^{-1}$]. Analytical HPLC analysis was performed on an HP Agilent 1100series HPLC system using an Agilent Poroshell 120 EC-$C_{18}$ (150 mm × 2.10 mm, 4 $\mu$m) column at 40 °C oven temperature and a detection wavelength of 254 nm. The mobile phase consisted of mixtures of acetonitrile, Milli-Q-grade water, and 10% of a 0.1% solution of formic acid in Milli-Q-grade water (injection volume = 5−20 $\mu$L; flow rate = 0.5−0.7 mL $min^{-1}$). Electrospray ionization (ESI) mass spectra were recorded on an Agilent1100 series LC/MSD ion trap spectrometer in the positive ion mode (drying gas temperature = 350 °C, nebulizer pressure = 70 psi, capillary voltage = 3500 V, drying gas ($N_2$) = 12 L $min^{-1}$). The purity of the final compounds was confirmed by HPLC analysis and was higher than 95% in all cases.

#### 2,6-Dichlorobenzaldoxime (*2*)

*Procedure adapted from McIntosh et al.*

McIntosh, M.L., Naffziger, M.R., Ashburn, B.O., Zakharov, L.N., Carter, R.G., 2012. Highly regioselective nitrile oxide dipolar cycloadditions with ortho-nitrophenyl alkynes. Org Biomol Chem 10, 9204–9213. https://doi.org/10.1039/c2ob26267c

NaOH solution (6 M, 11.7 mL, 70.00 mmol, 2.45 equiv.) was added dropwise to a suspension of 2,6-dichlorobenzaldehyde (***1***, 5.00 g, 28.57 mmol, 1.00 equiv.) and hydroxylamine hydrochloride (2.98 g, 42.85 mmol, 1.50 equiv.) in EtOH/$H_2O$ (40 mL, 1:1 v/v) at 0 °C. After stirring at 0 °C for 2 h, an aqueous HCl solution (1 M, 20 mL) was added. The formed precipitate was filtered off, washed with water, and dried under reduced pressure to yield the desired product as a colorless powder (5.08 g, 26.74 mmol, 94%). $^{1}$H NMR (300 MHz, DMSO-$d_6$): $\delta$/ppm = 11.79 (s, 1H, NO*H*), 8.21 (s, 1H, C<u>*H*</u>(NOH)), 7.57–7.48 (m, 2H, $H$-$3^{Ph}$ und $H$-$5^{Ph}$), 7.40 (dd, $J$ = 9.0, 7.1 Hz, 1H, $H$-$4^{Ph}$). $^{13}$C NMR (75.7 MHz, DMSO-$d_6$): $\delta$/ppm = 143.8 (*C*H(NOH)), 133.9 ($C$-$2^{Ph}$ und $C$-$6^{Ph}$), 130.9 ($C$-$4^{Ph}$), 129.4 ($C$-$1^{Ph}$), 128.9 ($C$-$3^{Ph}$ und $C$-$5^{Ph}$). IR (ATR): $\bar{\nu}$ / $cm^{-1}$ = 3279, 1583, 1559, 1438, 1418, 1299, 1187, 1095, 982, 918, 879, 777, 770, 717, 679. mp: 144–147 °C.

### ML2-SA1 (*3)*

*Procedure adapted from Plesch et al.*

Plesch, E., Chen, C.-C., Butz, E., Scotto Rosato, A., Krogsaeter, E.K., Yinan, H., Bartel, K., Keller, M., Robaa, D., Teupser, D., Holdt, L.M., Vollmar, A.M., Sippl, W., Puertollano, R., Medina, D., Biel, M., Wahl-Schott, C., Bracher, F., Grimm, C., 2018. Selective agonist of TRPML2 reveals direct role in chemokine release from innate immune cells. Elife 7. https://doi.org/10.7554/eLife.39720

(Bis(trifluoroacetoxy)iodo)-benzene (PIFA, 2.72 g, 6.32 mmol, 1.20 equiv.) was added dropwise to a solution of 2,6-dichlorobenzaldoxime (***2***, 1.00 g, 5.26 mmol, 1.00 equiv.) and norbornene (743 mg, 7.89 mmol, 1.50 equiv.) in MeOH/$H_2O$ (5:1 v/v). After stirring at room temperature overnight, brine (50 mL) was added and extracted with ethyl acetate (3 × 100 mL). The combined organic extracts were dried over $Na_2SO_4$, filtered, and the solvent was removed under reduced pressure at 40 °C. After purification by column chromatography on silica gel (cyclohexane/ethyl acetate = 40:1 v/v), the desired product was obtained as colorless crystals (1.06 g, 3.76 mmol, 71%). $^1$H NMR (300 MHz, $CDCl_3$): $\delta$/ppm = 7.52–7.20 (m, 3H, $H$-3$'^{Ph}$–$H$-5$'^{Ph}$), 4.74 (d, $J$ = 8.4 Hz, 1H, $H$-7a), 3.54 (d, $J$ = 8.4 Hz, 1H, $H$-3a), 2.69 (d, $J$ = 4.5 Hz, 1H, $H$-7), 2.35 (d, $J$ = 3.8 Hz, 1H, $H$-4), 1.98 (d, $J$ = 10.6 Hz, 1H, $H$a-8), 1.69–1.43 (m, 2H, $H$a-5 und $H$a-6), 1.31 (d, $J$ = 10.6 Hz, 1H, $H$b-8), 1.25–1.07 (m, 2H, $H$b-5 und $H$b-6). $^{13}$C NMR (75.5 MHz, $CDCl_3$): $\delta$/ppm = 154.7 ($C$-3), 135.3 ($C$-2$'^{Ph}$ und $C$-6$'^{Ph}$), 130.9 ($C$-4$'^{Ph}$), 129.2 ($C$-1$'^{Ph}$), 128.4 ($C$-3$'^{Ph}$ und $C$-5$'^{Ph}$), 89.0 ($C$-7a), 60.0 ($C$-3a), 42.6 ($C$-7), 39.1 ($C$-4), 33.1 ($C$-8), 27.4 ($C$-5), 23.0 ($C$-6). IR (ATR): $\tilde{\nu}$ / cm$^{-1}$ = 2957, 2933, 1581, 1557, 1431, 1327, 1313, 1194, 1159, 1109, 980, 920, 873, 866, 784, 727. mp: 114–115 °C. ESI-MS: $m/z$ = 282.1 (100%, $[M+H]^+$)/284.1 (68%, $[M+H]^+$); $m/z$ calc. for $[C_{14}H_{13}Cl_2NO+H]^+$ ($[M+H]^+$): 282.0/284.0. Purity: >99% (HPLC, 254 nm, MeCN/$H_2O$ = 55:45 + 0.1% HCOOH, $t_R$ = 4.23 min).

**NMR Spectra**

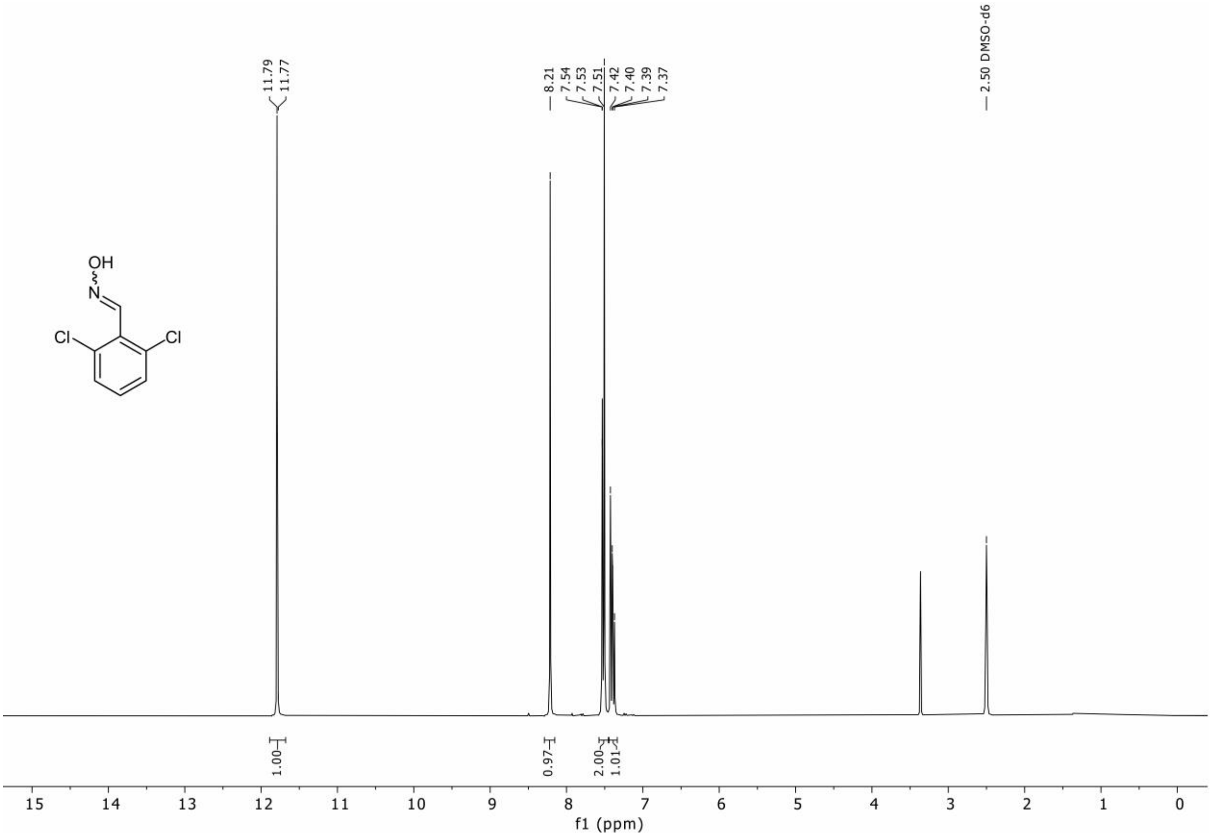


**Figure S1:** $^1$H NMR of 2,6-dichlorobenzaldoxime (***2***).

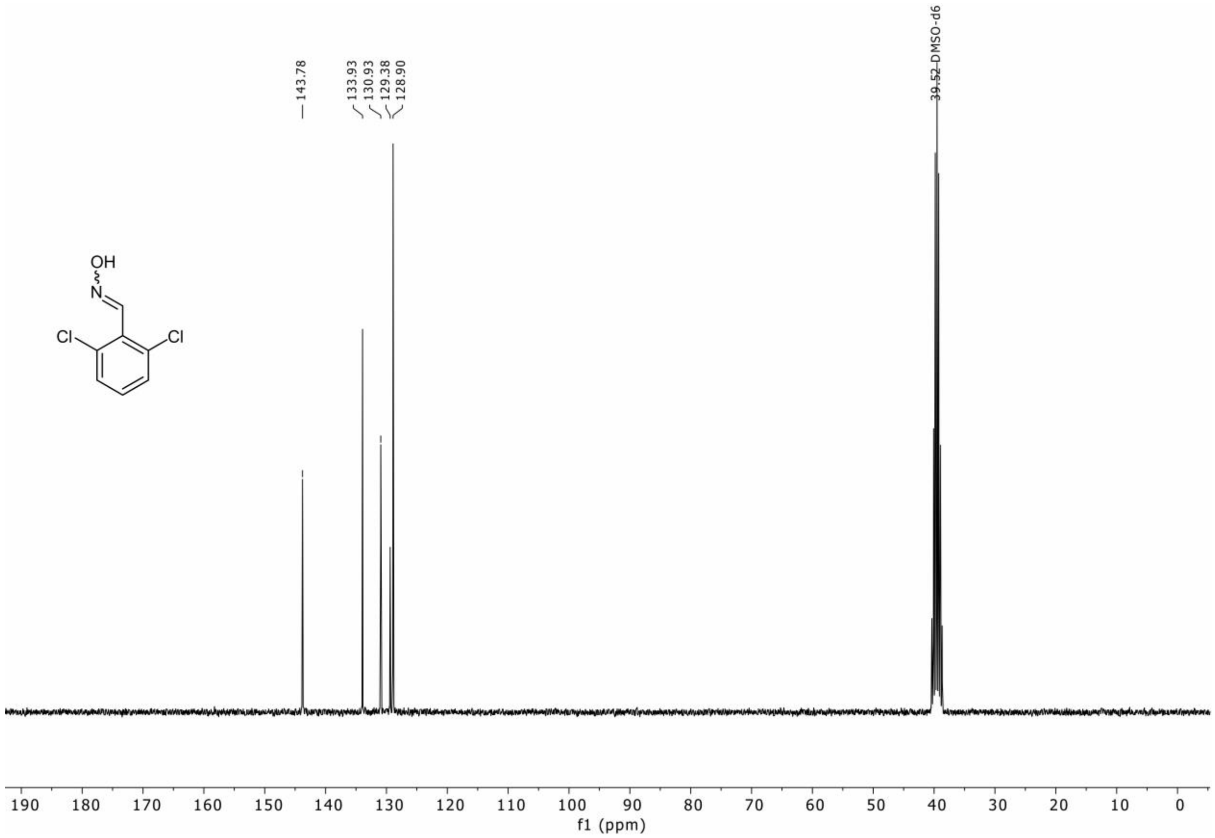


**Figure S2:** $^{13}C$ NMR of 2,6-dichlorobenzaldoxime (***2***).

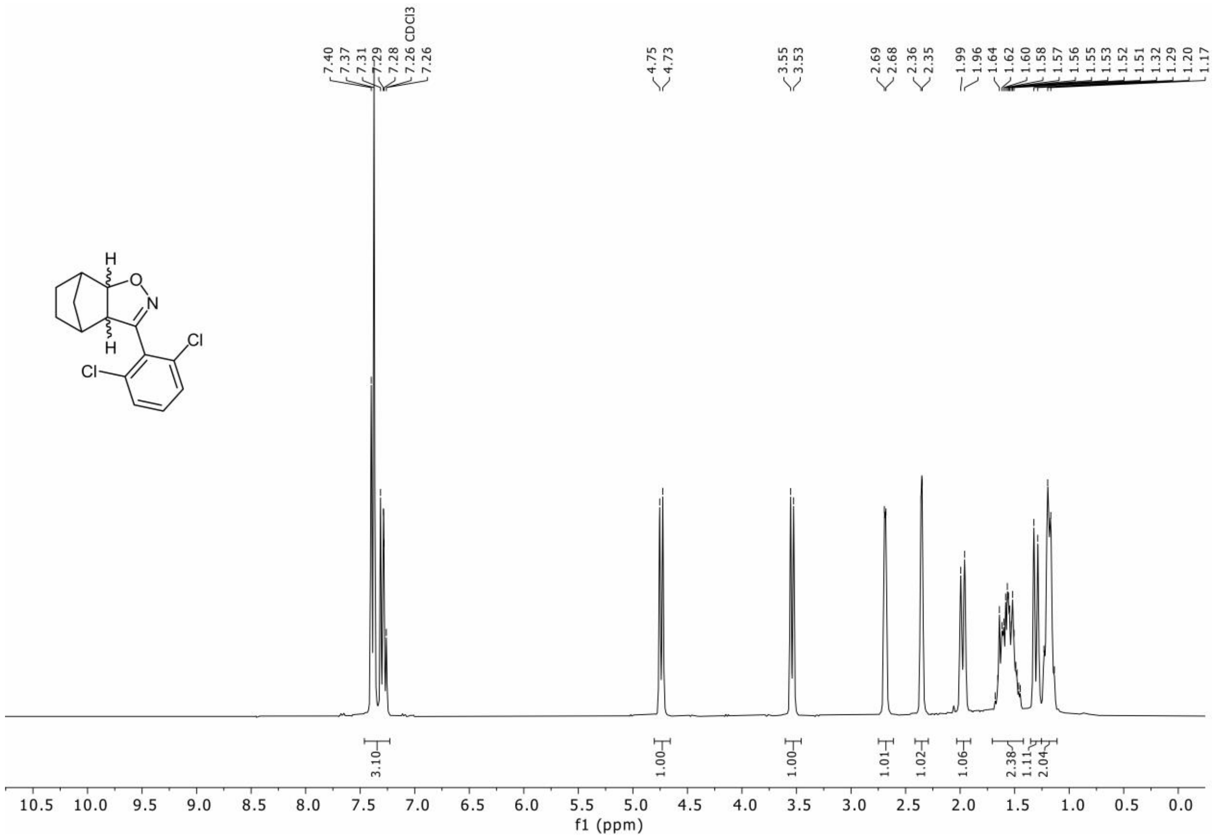


**Figure S3:** $^{1}$H NMR of ML2-SA1 (***3***).

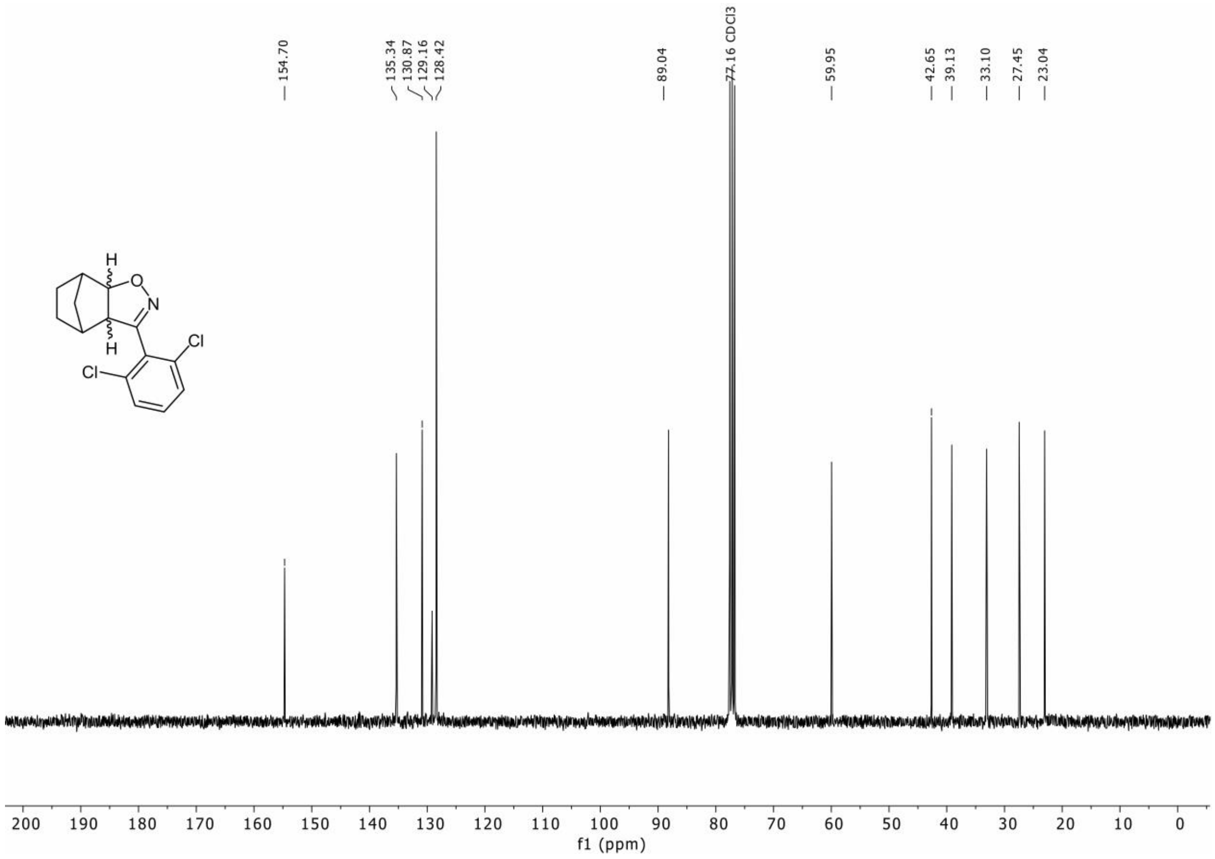


**Figure S4:** $^{13}C$ NMR of ML2-SA1 (***3***).

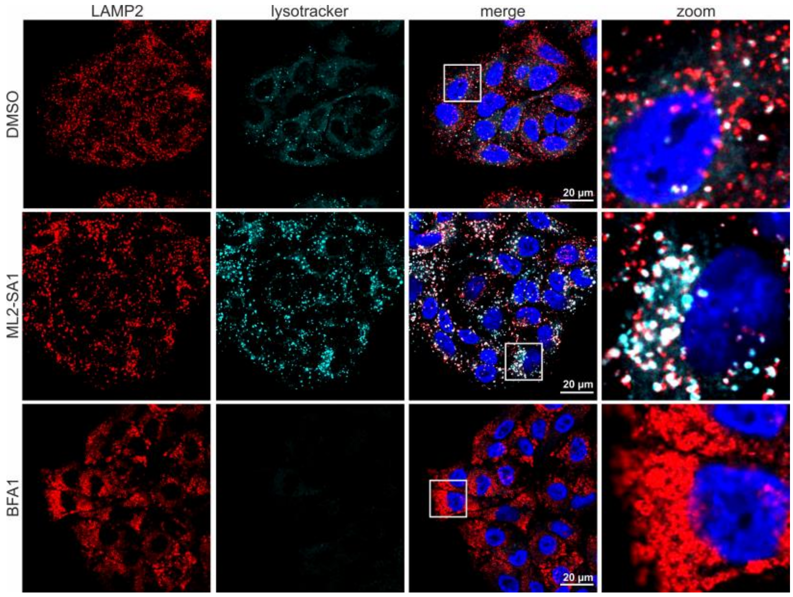


**Figure S5: Treatment of A549 cells with the TRPML2 agonist ML2-SA1 leads to increased acidification of the endolysosomal system.** Representative confocal laser scanning microscopy (cLSM) images of uninfected A549 cells treated with 100 µM ML2-SA1 and 50 nM BFA1, a V-type ATPase inhibitor. Cells were fixed 24 h post treatment and lysotracker (cyan) was added one hour before fixation to stain acidic organelles. Nuclei were stained with DAPI in blue and the lysosomal marker protein LAMP2 (red) was visualized using a LAMP2-specific antibody. Scale bar: 20 µm.

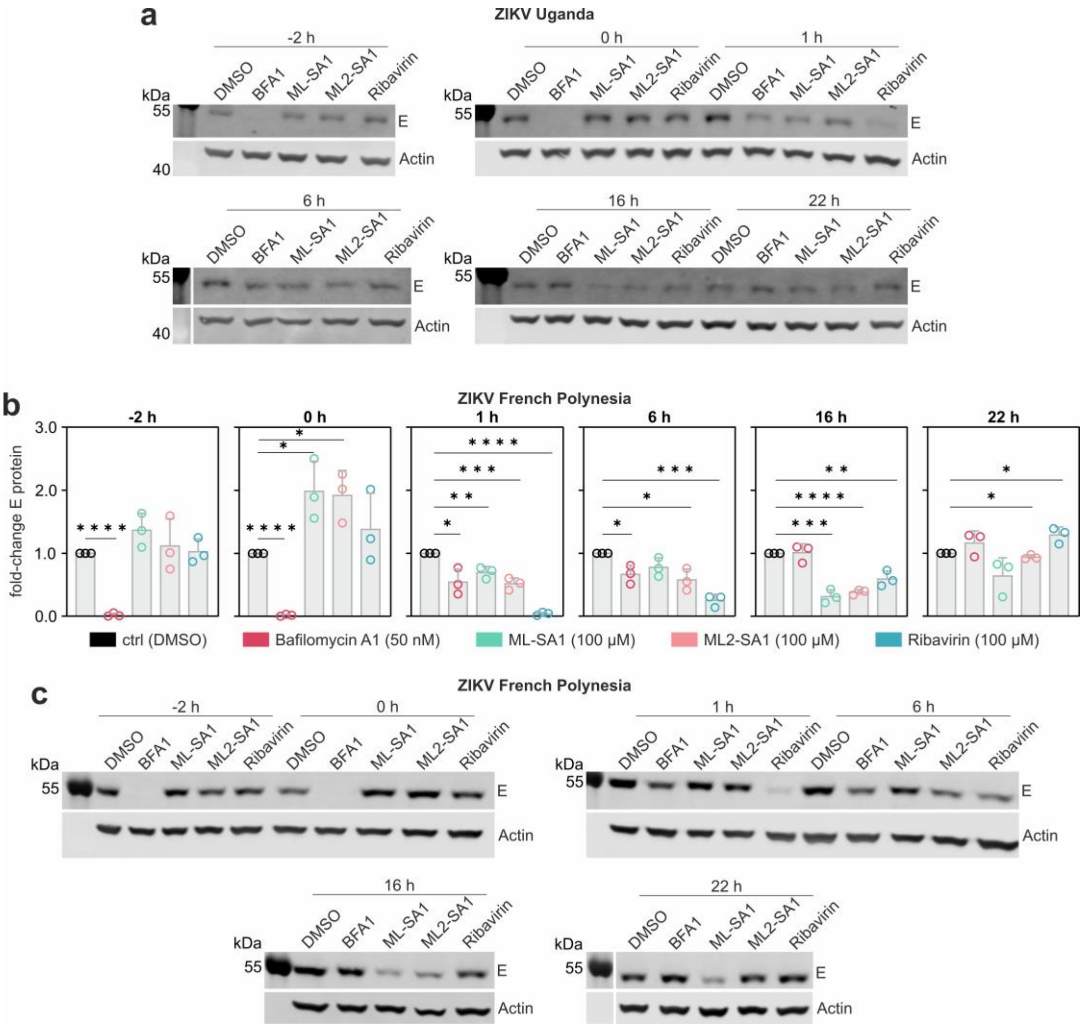


**Figure S6: ML2-SA1 affects late stages of the viral life cycle in A549 cells infected with ZIKV Uganda or ZIKV French Polynesia.** **(a)** Representative Western Blots of A549 cells infected with the ZIKV Uganda (976 Uganda) isolate for one hour (MOI = 1) prior to removal of unbound virus. Treatment with 50 nM Bafilomycin A1, 100 µM ML-SA1, 100 µM ML2-SA1 or 100 µM Ribavirin was started at −2h before infection, during infection, as well as 1 h, 6 h, 16 h and 22 h after infection (for timeline and relative fold-change of ZIKV Uganda E protein see Fig. 4a, main manuscript). **(b)** Relative fold-change of ZIKV E protein in A549 cells infected with ZIKV French Polynesia (H/PF/2013) treated with 50 nM Bafilomycin A1, 100 µM ML-SA1, 100 µM ML2-SA1 or 100 µM Ribavirin was assessed via Western Blotting (shown in c). Treatment was carried out as described above for ZIKV Uganda. **(c)** Representative Western Blots for cells infected with ZIKV French Polynesia (see b). Results were normalized to the respective DMSO control of each time point and are expressed as mean ± SD from $n = 3$ biological replicates. Statistical significance was determined by using an unpaired *t*-test. *$p < 0.05$, **$p < 0.01$, ***$p < 0.001$, ****$p < 0.0001$.

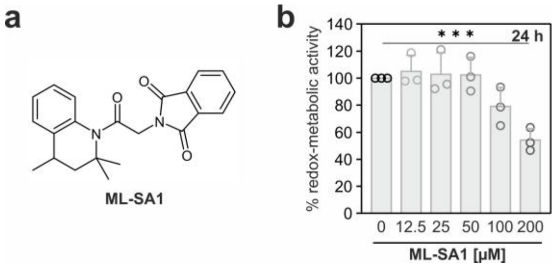


**Figure S7: The unspecific TRPML agonist ML-SA1 displays only mild cytotoxicity in A549 cells. (a)** Structure of ML-SA1. **(b)** Cell viability of A549 cells upon 24 h treatment with 12.5 µM to 200 µM ML2-SA1 determined with the PrestoBlue® assay. Values are expressed as % of intact cells referred to the experimental control. Data are normalized to the respective DMSO control and are expressed as mean ± SD from $n$ = 3 biological replicates, a representative blot is shown. Statistical significance was determined by using an unpaired $t$-test. *$p < 0.05$, **$p < 0.01$, ***$p < 0.001$, ****$p < 0.0001$.
